# Value-Based Evidence Accumulation as a Transdiagnostic Marker of General Distress

**DOI:** 10.64898/2026.02.16.706202

**Authors:** Helen Pushkarskaya, Catherine Maya Russell, Kenny Cheng, Jingzhu Chen, Christopher Pittenger

## Abstract

General distress cuts across psychiatric symptom domains, yet its computational correlates remain poorly defined. We examined whether drift rate—a core parameter indexing the efficiency of evidence accumulation—is more strongly associated with general distress than with domain-specific symptoms. In a cross-sectional online sample of 441 adults from the general population, participants completed a perceptual and value-based decision-making task, symptom assessments, and cognitive testing. Drift rates were estimated using hierarchical drift-diffusion modeling. Individuals with severe symptom elevations showed robust reductions in drift rate, particularly for value-based decisions. Mixed-effects models demonstrated that general distress, indexed by the Positive Symptom Distress Index, was more strongly associated with value-based than perceptual drift rate, even after accounting for all symptom domains. Value-based drift rate also explained variance in general distress beyond that accounted for by elevated symptoms across domains and selectively attenuated associations with somatization and paranoid symptoms. These findings suggest that value-based evidence accumulation captures a transdiagnostic component of distress-related impairment that is not reducible to symptom burden alone.

## Introduction

Psychiatric symptom severity and subjective distress are only partially aligned across conditions. Individuals with comparable symptom profiles often differ markedly in perceived burden, functional interference, and need for clinical care. Identifying computational processes that track this dissociation may help clarify mechanisms underlying generalized distress beyond diagnostic categories.

Drift rate, derived from sequential sampling models, indexes the efficiency with which noisy information is accumulated toward a decision. Prior work links reduced drift rate to psychopathology across tasks and diagnoses, but it remains unclear whether such reductions reflect domain-specific symptoms or a more general dimension of distress-related impairment. Importantly, evidence accumulation during value-based decisions may be particularly sensitive to affective and motivational disturbances that characterize generalized distress.

Here, we tested whether drift rate is more strongly associated with general distress than with domain-specific symptom dimensions, and whether this relationship differs between perceptual and value-based decisions. We further examined whether drift rate explains variance in distress beyond that attributable to elevated symptoms across domains.

## Methods

### Participants

All procedures were approved by Yale Institutional Review Board (IRB). Participants were recruited via Amazon Mechanical Turk and provided consent in accordance with IRB regulations and guidelines. Participants were prescreened for data quality and task compliance using attention checks, response-consistency items, and the MMPI Lie subscale. Only participants whose responses to pre-screen questions (e.g., attention check questions, scores on the Lie scale greater than 7) passed rigorous quality control were enrolled. The final sample included 213 men and 222 women (ages 18–67).

### Measures

Participants completed the Perceptual and Value-Based Decision-Making (PVDM) task, a computerized adaptive measure of cognitive ability (KBIT-2 Matrices), the Brief Symptom Inventory (BSI), and demographic questionnaires.

#### Perceptual and value-based decision making (PVDM) task. ^**1, 2**^

The task interleaves perceptual (PDM) and preferential/value-based (VDM) decision making across four sequential phases to dissociate information accumulation from value-to-action translation. In Phase I (Rating), participants provide individual perceptual and value ratings for 80 emotionally neutral grayscale images presented one at a time in pseudorandom order. Perceptual ratings estimate the percentage of black ink in each image (10–90% in 10% increments), while value ratings assess liking on a 9-point scale. In Phase II (Choice), an adaptive algorithm uses these ratings to generate individualized PDM and VDM choice sets with four levels of difficulty, defined by objective differences in image darkness for PDM (10%, 20%, 40%, or 50%) and by differences in individual liking ratings for VDM (1, 2, 4, or 5 points; Figure 1C). Participants then complete separate PDM and VDM choice blocks (52 trials each), selecting either the darker image (PDM) or the preferred image (VDM); choice accuracy is defined objectively for PDM and relative to each participant’s own ratings for VDM. In Phase III, participants repeat value ratings for images included in the choice blocks to assess rating stability. In Phase IV, PDM and VDM blocks are repeated with randomized trial order, and 25% of trials are followed by brief negative feedback (“The choice is wrong” for PDM; “The choice is inconsistent” for VDM), allowing examination of how experimentally induced negative affect modulates information processing and decision dynamics. Participants are given up to 6 seconds to respond on each trial, though most responses occur within ∼2 seconds. Average time to complete the task based on over 600 participants is ∼25 min.

**Figure 1.**
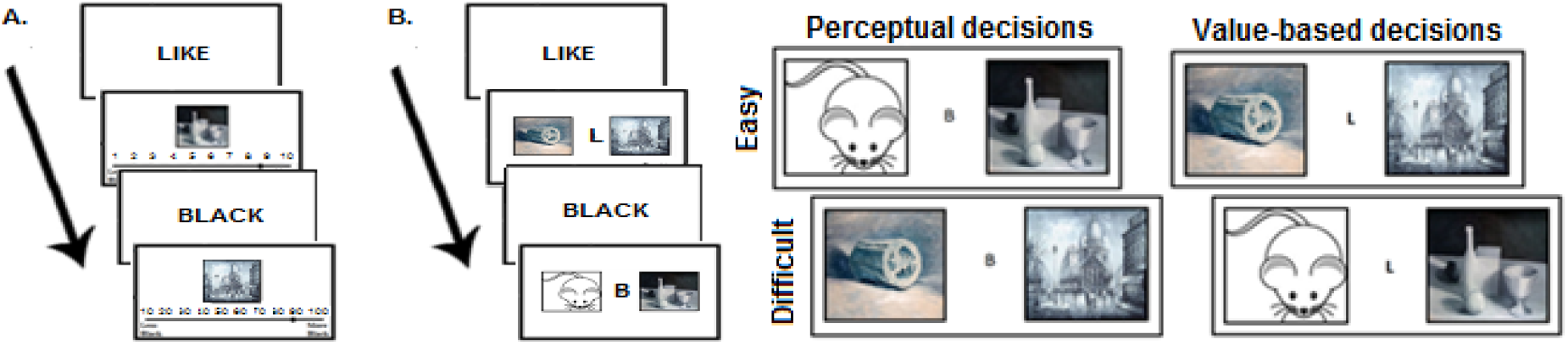
Perceptual and Value Based Decision Making. **A.** Rating Blocks. **B** Choice Blocks. **C**. Sample Stimuli.

#### The Brief Symptom Inventory (BSI)^3^

is a 53-item self-report measure designed to assess current psychological symptom burden across nine primary symptom dimensions (Somatization, Obsessive–Compulsive, Interpersonal Sensitivity, Depression, Anxiety, Hostility, Phobic Anxiety, Paranoid Ideation, and Psychoticism). Respondents rate the degree to which they have been distressed by each symptom over the past week on a 5-point Likert scale ranging from 0 (“not at all”) to 4 (“extremely”). In addition to domain scores, the BSI provides global indices of distress, including the Global Severity Index (GSI), Positive Symptom Total (PST), and Positive Symptom Distress Index (PSD), which capture complementary aspects of overall symptom burden and intensity. Raw scores can be converted to sex-normed T scores based on community and clinical reference samples. The BSI demonstrates good internal consistency, test–retest reliability, and convergent validity across community and clinical populations, and is widely used in psychiatric and epidemiological research (Cronbach’s α = 0.90-0.95).

Raw BSI scores were converted to sex-normed T scores. Provisionally elevated symptoms were defined as T > 63, and severely elevated symptoms as T > 75. General distress was indexed using the BSI Positive Symptom Distress Index (PSD), which reflects average symptom intensity independent of symptom count.

### Computational Modeling

Response time (RT) and accuracy are recorded on each of 208 trials to be analyzed using hierarchical Bayesian parameter estimation in the Drift Diffusion Model (HDDM).^4^ Trials with RT < 0.2 s and extreme outliers (RT < mean—3 SD or RT > mean + 3 SD) were discarded; the minimum number of remaining trials across participants was 180. HDDM allows simultaneously estimating the group level and the individual level parameters for each of the four conditions of the task. Our sample of 441 participants with at least 180 data points per participant was well powered for the planned analysis.^5^ Drift rates were estimated separately for perceptual and value-based trials and under neutral and negative affect conditions. Analyses focused on condition-specific drift-rate estimates.

### Statistical Analysis

Group differences in drift rate were examined using clinically meaningful symptom thresholds (T < 63 vs. T > 75). To dissociate general distress from symptom-specific effects, hierarchical mixed-effects models were fit including age, IQ, task condition, and all symptom domains residualized with respect to PSD. Incremental predictive value of drift rate was assessed via nested regression models predicting PSD.

## Results

### Drift Rate and Severe Symptom Elevation

Individuals with very severe symptoms (T > 75) exhibited robust reductions in drift rate (Figure 2). Paranoid and somatization symptoms were associated with reduced drift rates in both perceptual and value-based decisions, with larger effects in value-based trials (d ≈ ™0.32 to ™0.40). Obsessive–compulsive, hostility, anxiety, and interpersonal sensitivity symptoms were associated with significant reductions selectively in value-based drift rate (d ≈ ™0.15 to ™0.25). Depressive, phobic, and psychotic symptoms showed no reliable associations with drift rate.

**Figure 2.**
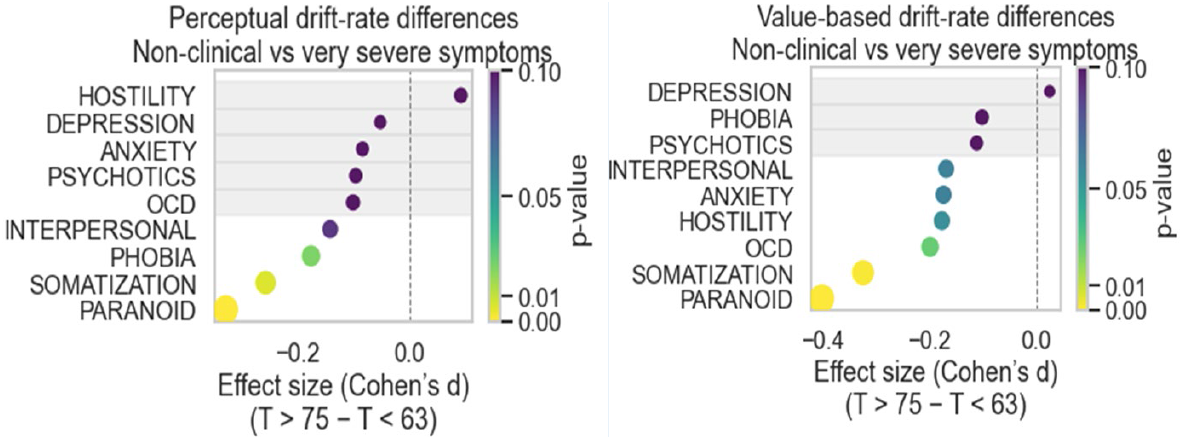
Drift-rate differences between non-symptomatic and very severe symptom groups across conditions. Effect sizes (Cohen’s *d*) for drift-rate differences between non-clinical (T < 63) and very severe (T > 75) symptom groups are shown separately for perceptual (top) and value-based (bottom) conditions. Negative values indicate lower drift rates in the very severe group. Point color reflects raw *p*-values, and gray shading denotes non-significant effects (*p* > 0.10). Dashed vertical lines indicate no group difference.

In mixed-effects models accounting for demographics, cognition, task condition, and all symptom domains, adding general distress (PSD) significantly improved model fit. Drift-rate associations with PSD were stronger for value-based than perceptual decisions (PSD main effect: β = ™0.063, p = 0.0035; PSD × condition interaction: β = 0.09, p = 5 × 10^−6^; Figure 3A). Among residualized symptom domains, only paranoid tendencies remained independently associated with drift rate (Figure 3B).

**Figure 3.**
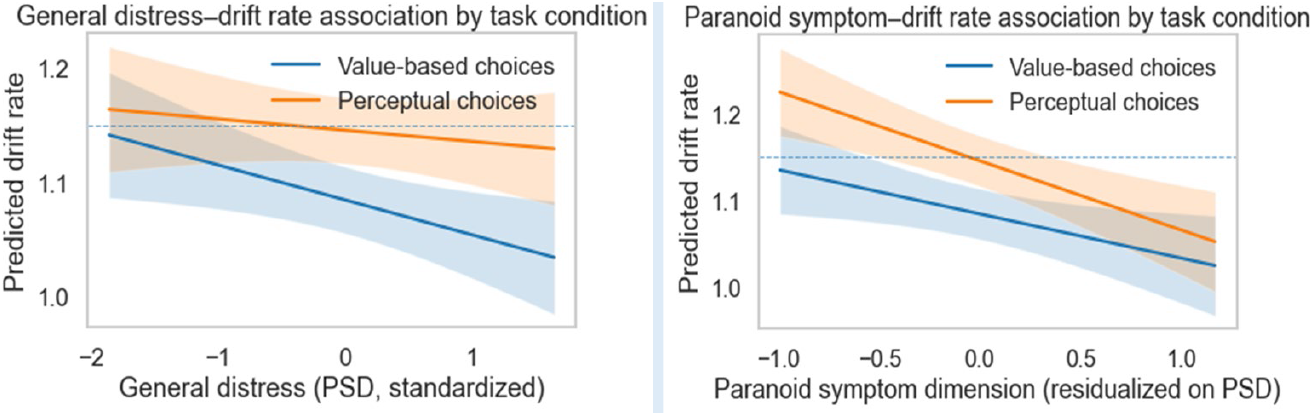
Drift-rate associations with general distress and paranoid symptoms by condition. Predicted drift rate is shown as a function of standardized general distress (left) and paranoid symptom severity residualized on distress (right) for value-based and perceptual trials. Shaded bands indicate 95% confidence intervals. Associations are stronger for value-based than perceptual decisions.

### Incremental Prediction of Distress

Including value-based drift rate significantly improved prediction of generalized distress beyond that explained by elevated symptoms across domains (ΔR^2^ ≈ .01; p ≈ 0.01). Perceptual drift rate did not contribute additional explanatory power. Inclusion of value-based drift rate attenuated associations between distress and somatization (β reduction ∼25–30%) and paranoid symptoms (β reduction ∼30–40%), with some effects losing statistical significance. Obsessive–compulsive symptoms showed weaker attenuation (β reduction ∼10–15%) and remained marginally significant (p = 0.06).

## Discussion

Across a large community sample, reduced drift rate—particularly during value-based decisions—was more closely linked to generalized distress than to most symptom-specific dimensions. These associations persisted after controlling for cognitive ability, demographics, and symptom burden, and value-based drift rate explained unique variance in distress beyond elevated symptoms.

These findings support value-based drift rate as a computational marker of generalized distress with potential relevance for transdiagnostic assessment. Longitudinal and clinical samples will be needed to determine whether this marker predicts functional outcomes or treatment response.

## Data Availability

De-identified data and analysis code will be made available upon reasonable request.

